# Phylogenetic analysis reveals unexplored fungal diversity on skin

**DOI:** 10.1101/2023.11.22.568368

**Authors:** Pedro Nicolas Brito, Luciana Campos Paulino

## Abstract

Despite recent advances, to date there is not full knowledge of microbial diversity inhabiting various organs of human body. Skin harbors a complex microbiome that might affect our health positively and negatively. Fungal communities from skin are dominated by *Malassezia* yeasts. Traditionally, they were thought to be causative agents of skin diseases; however, their role is controversial, and the possible implication of specific species and subtypes remains unclear. Previously, we have conducted two fungal community surveys in healthy skin and dandruff/sebohrreic dermatitis, and have detected prevalent *Malassezia* organisms that could not be assigned to any known species. The usage of distinct ITS rDNA regions did not allow sequence comparison between studies. Here we report molecular characterization and phylogenetic analysis of unidentified *Malassezia* organisms, aiming to increase knowledge in fungal microbiome from skin. Findings suggested that a highly prevalent organism might belong to a novel *Malassezia* species. Results also revealed uncertain taxonomic assignments, even in the case of accepted species. Correct assignment of species and intraspecific variants is relevant considering that specific taxa might be directly involved in disease development. Despite high prevalence, organisms might have remained undiscovered due to difficulties in culturing *Malassezia*. Challenges and future perspectives for skin fungal microbiome studies are discussed. We address issues to be overcome for unraveling the complete skin microbial diversity and its relation to health and disease.

---

Despite recent advances, to date there is not full knowledge of microbial diversity inhabiting various organs of human body^1^. Skin microbiome protects against pathogens, synthesizes essential compounds and modulates immune response. However, unbalances in skin microbial communities might be related to skin diseases^2^. Fungal communities inhabiting skin are dominated by *Malassezia* yeasts^3^. Traditionally, these lipophilic yeasts had been pointed out as causative agents of inflammatory skin diseases, and have been detected in psoriasis, atopic dermatitis, pityriasis versicolor, dandruff and seborrheic dermatitis^4^. Nevertheless, their possible role in disease development is unclear^5^, as well as the implication of specific species and subtypes.

*Malassezia* genus comprises fastidious organisms and media requirements differ among species^4^, representing a challenge for isolation, preservation, characterization, and identification of species. Currently there are 18 formally described species, 9 usually found in humans^6^.

Previously, we have characterized skin *Malassezia* communities from seborrheic dermatitis and healthy subjects, using RFLP and 5.8S/ITS2 rDNA sequencing^7^. We have detected intraspecific variation, and three phylotypes that were not assigned into any known species, including highly prevalent ones. Two phylotypes have been reported before in health and psoriasis^8^, although had not been characterized, and a new phylotype has been reported for the first time (phylotype 5).

Subsequently, we have investigated bacterial and fungal skin communities in health and dandruff through ITS1 Next Generation Sequencing^9^. In addition to genus level analyses, we built a database for studying *Malassezia* at species level, and found four uncharacterized organisms. Subgroups 1 and 2 were detected in high proportions in most samples. Because the studies had distinct goals, different ITS regions were sequenced, thus we were unable to compare sequences and determine whether they belong to the same organisms.

Characterizing potential novel species, particularly prevalent ones, is important for establishing relationships with skin diseases. Therefore, we carried out molecular characterization and phylogenetic analysis of potential novel *Malassezia* species from skin.

Research protocol was approved by UFABC Institutional Review Boards (CAAE 41835815.4.0000.5594) and was conducted according to the principles expressed in the World Medical Association Declaration of Helsinki. All subjects provided written informed Consent prior to any study-related procedures. Samples from healthy and dandruff/seborrheic dermatitis skin^7,9^ were screened based on proportions of uncharacterized *Malassezia*. A dandruff scalp sample (over 95% of subgroup 1), and a healthy forehead sample (over 75% of subgroup 2) were selected^9^.

PCR primers were combined to allow amplification of both ITS rDNA regions analyzed previously (Fig. 1). PCR products were cloned, inserts were screened by length in electrophoresis gels, and Sanger sequenced. Uncharacterized *Malassezia* sequences were compared to Genbank database using BLAST algorithm, as well as to sequences from previously detected uncharacterized organisms. This approach enabled us to compare organisms detected in previous studies targeting different DNA regions. Results suggested that ITS2-based phylotype 5 and ITS1-based subgroup 1 correspond to the same organism. Moreover, BLAST analysis of complete ITS sequence from subgroup 1 did not support its assignment into any formally described species. Conversely, subgroup 2 complete ITS sequence analysis revealed high identity with *Malassezia restricta*.

**Fig. 1.**
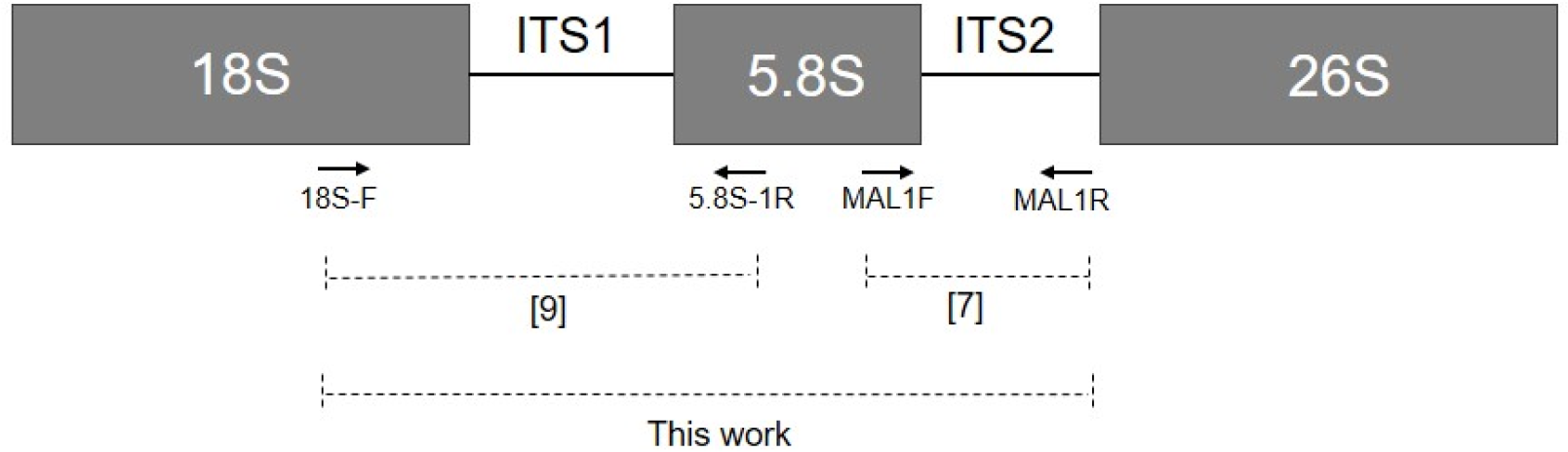
Schematic representation of fungal ribosomal operon, showing primer annealing sites (arrows). 18SF/5.8S–1R and Mal1F/Mal1R amplicons have been reported by references^9^ and^7^, respectively. Estimated length of 18SF/Mal1R amplicon: 550 to700 bp.

Phylogenetic analyses were performed using 5.8S/ITS2 sequence portion, since ITS1 indels do not allow accurate alignment. Newly obtained sequences from subgroups 1 and 2 were included, as well as previously reported uncharacterized phylotypes, *Malassezia* type strains, and additional *M. restricta* strains for better clade resolution (Fig. 2).

**Fig. 2.**
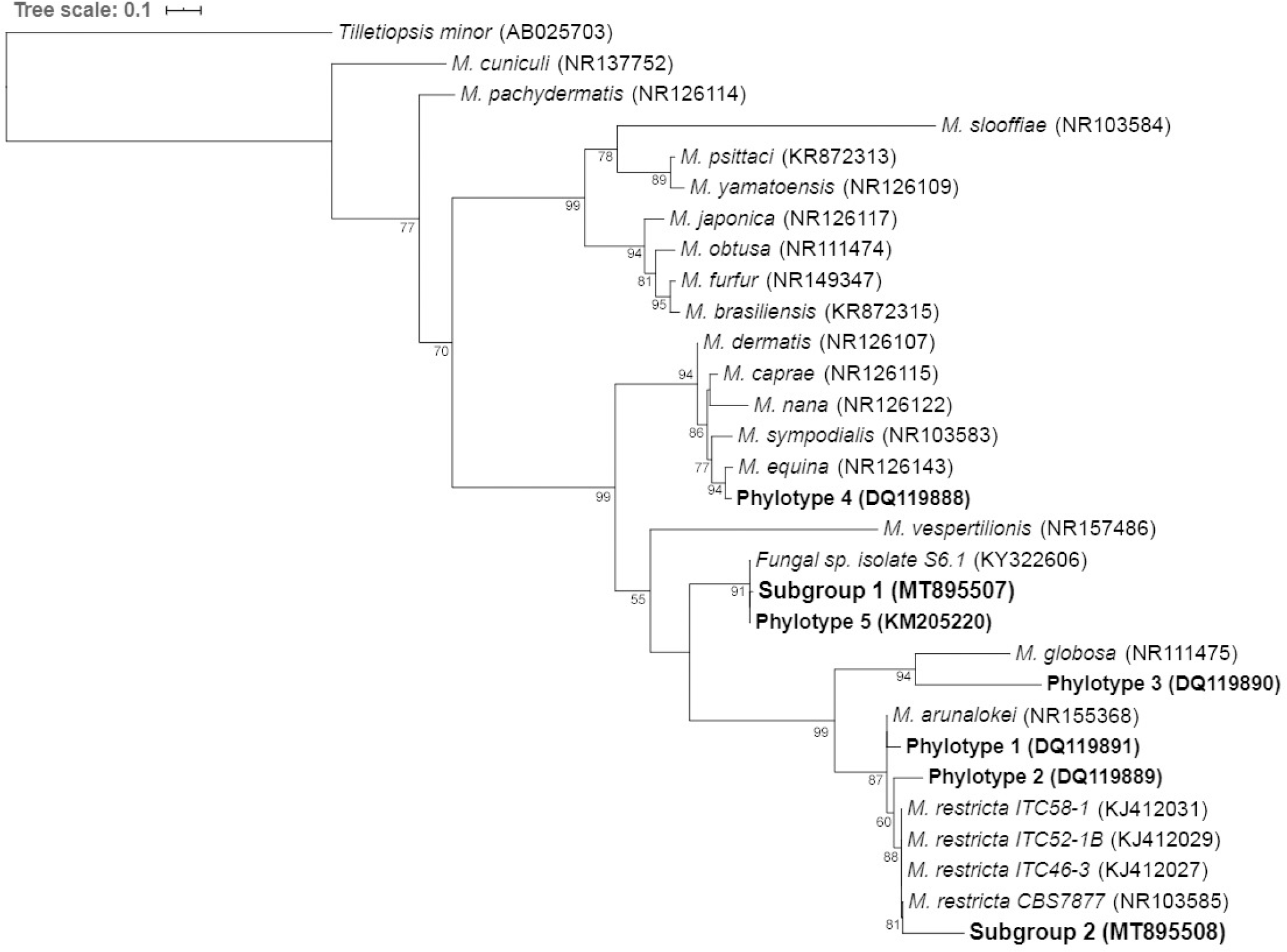
Molecular phylogenetic tree inferred from Maximum Likelihood analysis of 5.8S/ITS2 sequences from *Malassezia* sp., including uncharacterized organisms (in bold type). Branch support aLRT (Approximate Likelihood Ratio Test) values above 50% are shown. GenBank accession numbers are indicated between parentheses.

Subgroup 1/phylopype 5 sequences did not cluster together with any described *Malassezia* species, supporting the novel species hypothesis. In contrast, subgroup 2 was closely related to *M. restricta*. Phylotypes 1 and 2 belong to the same clade, along with *M. arunalokei*. The corrected species assignment depends on target DNA region and database sequence availability, which might help explaining why previous works failed in assigning the organisms to *M. restricta*.

Subgroup 2 branch length suggests it might be a variant of *M. resctricta*. Relatedness to *M. arunalokei*, together with high degree of *M. restricta* intraspecific variation previously reported^7^, suggests that *M. restricta/M. arunalokei* could comprise a species complex.

Furthermore, findings indicated that phylotyes 3 and 4 clustered together with *M. globosa* and *M. equina*, respectively. Nevertheless, it is unclear whether they should be assigned to these species, as phylogenetic distances are similar or greater than distances between some of the taxonomic species (Fig. 2).

Correctly assigning taxonomic species and intraspecific variants is particularly important considering that diseases might be associated with *Malassezia* subtypes^5^. Despite advantages of molecular-based methods, culture approaches are useful for accessing many biological features, as well as for organism preservation. Moreover, their cost-effectiveness makes them the only feasible option depending on the local socioeconomic situation. However, *Malassezia* culture can be challenging. Our attempts to isolate uncharacterized organisms from skin included different culture media, modified media composition, lipid supplementation, temperature and other growth conditions, but were not successful. Despite their prevalence, they might remain uncultured and therefore uncharacterized due to growth issues. Possibly, they are overgrown by less fastidious species, or require undetermined culture conditions. The establishment of effective culture conditions should be pursued, in order to contribute to skin fungal diversity knowledge and its possible role in skin diseases, as well as microbial interactions in health.

In conclusion, we presented evidences of prevalent novel *Malassezia* species associated with health and diseased skin, suggesting an undisclosed skin-associated microbial diversity. Molecular-based approaches should focus not only on extensive data, but also on increasing resolution to allow strain-level analyses. Further study expansion could also benefit from standardized protocols and assembly of public databases. In parallel, it is important to improve culture-based methods, as well as establishing human sample biobanks and culture collections.

Multi-country studies are necessary for accessing microbial variation due to genetic factors, socioeconomic status and cultural features, as well as addressing issues of particular interest of each country. Human microbiome research in Brazil has progressed recently^10^, but it requires more effective public policies for supporting individual research groups and local consortia initiatives. Finally, bridges should be constructed to connect scientific research and clinical practice, accelerating knowledge transfer and converting it into new possibilities for diagnostics, therapy, and prevention of diseases.

## Acknowledgments

The authors thank Juliana Tonini for technical support.

## Data Availability

Nucleotide sequences reported here are available in the GenBank database under the accession numbers MT895507 and MT895508.

